# CA2 Neuronal Activity Controls Hippocampal Oscillations and Social Behavior

**DOI:** 10.1101/190504

**Authors:** Georgia M. Alexander, Logan Y. Brown, Shannon Farris, Daniel Lustberg, Caroline Pantazis, Bernd Gloss, Nicholas W. Plummer, Natallia V. Riddick, Sheryl S. Moy, Patricia Jensen, Serena M. Dudek

## Abstract

Hippocampal oscillations arise from coordinated activity among distinct populations of neurons and are associated with cognitive functions and behaviors. Although much progress has been made toward identifying the relative contribution of specific neuronal populations in hippocampal oscillations, far less is known about the role of hippocampal area CA2, which is thought to support social aspects of episodic memory. Furthermore, the little existing evidence on the role of CA2 in oscillations has led to conflicting conclusions. Therefore, we sought to identify the specific contribution of CA2 pyramidal neurons to brain oscillations using a controlled experimental system. We used excitatory and inhibitory DREADDs in transgenic mice to acutely and reversibly manipulate CA2 pyramidal cell activity. Here, we report on the role of CA2 in hippocampal-prefrontal cortical network oscillations and social behavior. We found that excitation or inhibition of CA2 pyramidal cells bidirectionally regulated hippocampal and prefrontal cortical low gamma oscillations and inversely modulated hippocampal ripple oscillations. Further, CA2 inhibition impaired social approach behavior. These findings support a role for CA2 in low gamma generation and ripple modulation within the hippocampus and underscore the importance of CA2 neuronal activity in extrahippocampal oscillations and social behavior.

---

Area CA2 of the hippocampus has become appreciated as a distinct subfield of hippocampus based on several molecular, synaptic, anatomical, and functional properties (see^1^ for review). We and others have recently identified similarities and differences between CA2 and the neighboring CA1 and CA3 subfields based on action potential firing *in vivo^2,6^*. In addition to action potential firing, another form of neuronal communication is achieved through synchronized oscillations^7^, which reflect the summated electrical activity of a population of neurons and can be detected in local field potentials (LFPs). CA1 and CA3 networks propagate oscillations in three primary frequency bands: theta (~5-10 Hz), gamma (~30-100 Hz) and sharp-wave ripples (~100-250 Hz). A few studies have reported properties of network oscillations in CA2^6,8,9^, but none of them have examined CA2 gamma oscillations or the impact of CA2 oscillations on extrahippocampal structures.

In the hippocampus, high and low gamma oscillations are thought to arise from two distinct sources and likely play separate roles in memory. High gamma (~60-100 Hz) oscillations in CA1 are prevalent in stratum lacunosum-moleculare^11^, co-occur with high gamma oscillations in medial entorhinal cortex (MEC)^ıν^, and are impaired by lesioning of EC^12^, leading to the conclusion that high gamma oscillations arise from MEC. High gamma is thought to contribute to memory encoding because high gamma power is increased upon exploration of novel stimuli^13,14^. Low gamma (~30-55 Hz) oscillations in CA1 are prevalent in stratum radiatum^11^, synchronize with low gamma in CA3^15^, and become more evident upon EC lesioning^12^, supporting the conclusion that low gamma oscillations arise from CA3. Low gamma oscillations are believed to promote memory retrieval because the magnitude of low gamma coupling to theta oscillations correlates with performance on learned behavioral tasks^16,17^. Interestingly, complete silencing of the synaptic output of CA3 with tetanus toxin light chain does not completely impair low gamma oscillations^18^, suggesting the presence of another source of low gamma oscillations.

Another prominent oscillation seen in hippocampus is sharp-wave ripple oscillations, which are high frequency (~100-250 Hz), short-duration electrical events prominently seen in LFP recordings from CA1 during awake immobility and slow wave sleep^19^. Sharp waves are thought to arise from the synchronous firing of CA3 pyramidal cells, which depolarizes the apical dendrites of CA1 pyramidal cells. The synchronous CA3 firing recruits excitatory and inhibitory neurons in CA1 to generate ripples^19,20^. A role for CA2 neurons in sharp-wave ripples has recently been suggested based on three *in vivo* electrophysiology studies^6,8,9^, although consensus has not been reached on the precise role that these neurons play. Kay et al. found that CA2 is the only hippocampal subregion to have a substantial population of neurons that cease firing during ripples (termed ‘N cells’), whereas nearly all pyramidal cells queried in neighboring subfields fired during ripples. Although not associated with ripples, these N cells fired at high rates during low running speed or immobility^6^. Oliva et al. later reported that CA2 pyramidal cell activity ramps up before the onset of sharp-wave ripples, leading these authors to conclude that CA2 neurons play a leading role in ripple generation. By contrast, Boehringer et al. later found that chronic silencing of CA2 pyramidal cell output leads to the occurrence of epileptic discharges arising from CA3, which the authors suggested reflect anomalous ripple oscillations. Accordingly, findings of the Boehringer study do not appear to support the conclusion of Oliva et al. that CA2 neurons initiate ripples. Given the disparate conclusions of these reports, further study is required to clarify the role of CA2 neuronal activity in ripple generation.

Area CA2 has recently been recognized for its role in processing long term memories containing socially relevant information in rodents^2,21–23^. Interestingly, a mouse model of schizophrenia that shows hypoactive CA2 pyramidal cells *in vitro* also shows impaired social behavior^24^. Further, long range synchrony between hippocampus and prefrontal cortex (PFC), including low gamma coherence, is impaired in another mouse model of schizophrenia^25^, raising the question of how altering CA2 pyramidal cell activity experimentally may impact social behavior and synchrony between hippocampus and PFC.

In this study, we present evidence that selective, acute activation or inhibition of CA2 pyramidal cells using Cre-dependent expression of Gq- and Gi-coupled DREADD receptors (hM3Dq and hM4Di^26,27^, respectively) bidirectionally modulates low gamma oscillations in both hippocampus and PFC and ripple occurrence in hippocampus. Further, manipulation of CA2 with the inhibitory DREADD affected behavior in one measurement of social function.

## RESULTS

### Increasing CA2 pyramidal cell activity increases hippocampal and prefrontal cortical low gamma power

To gain selective genetic access to molecularly-defined CA2, we generated a tamoxifen-inducible mouse line, *Amigo2-icreERT2*. When combined with a Cre-dependent tdTomato reporter mouse line^28^, we observed robust expression of tdTomato in CA2 of brain sections from *Amigo2-icreERT2+;* ROSA-tdTomato+/-mice treated with tamoxifen. Expression of tdTomato colocalized with the CA2 pyramidal cell marker, PCP4^29^ (N=6 mice; Fig. S1), and a marker of hippocampal pyramidal neurons (N=6; Fig S1, S2) but not inhibitory neurons (N=3; Fig S1, S2). Expression of tdTomato was also observed in extra-hippocampal brain structures and associated with vasculature. In control experiments, *Amigo2-icreERT2+;* ROSA-tdTomato+/-animals treated with corn oil (the tamoxifen vehicle) showed no tdTomato expression (N=3; Fig. S3).

Infusion of AAVs encoding Cre-dependent hM3Dq (Fig. 1A-C, E) or hM4Di (Fig. 1D) with the neuron-specific human synapsin promoter into *Amigo2-icreERT2+* mice allowed for selective expression of mCherry-DREADD in CA2 pyramidal neurons without expression in fasciola cinerea, outside of the hippocampus, or in the vasculature, as detected by co-expression of mCherry with PCP4 (N=4; Fig 1A-B, E). Expression of mCherry also colocalized with the pyramidal cell marker, CaMKIIα (N=4; Fig. 1C, Fig. S2), but not the interneuron marker, glutamic acid decarboxylase (GAD) in GAD-eGFP+; *Amigo2*-icreERT2+ mice (N=4; Fig. 1D, Fig. S2). In control *Amigo2*-icreERT2-mice infused with hM3Dq AAV, mCherry expression was absent (N=4; Fig. S3).

**Figure 1.**
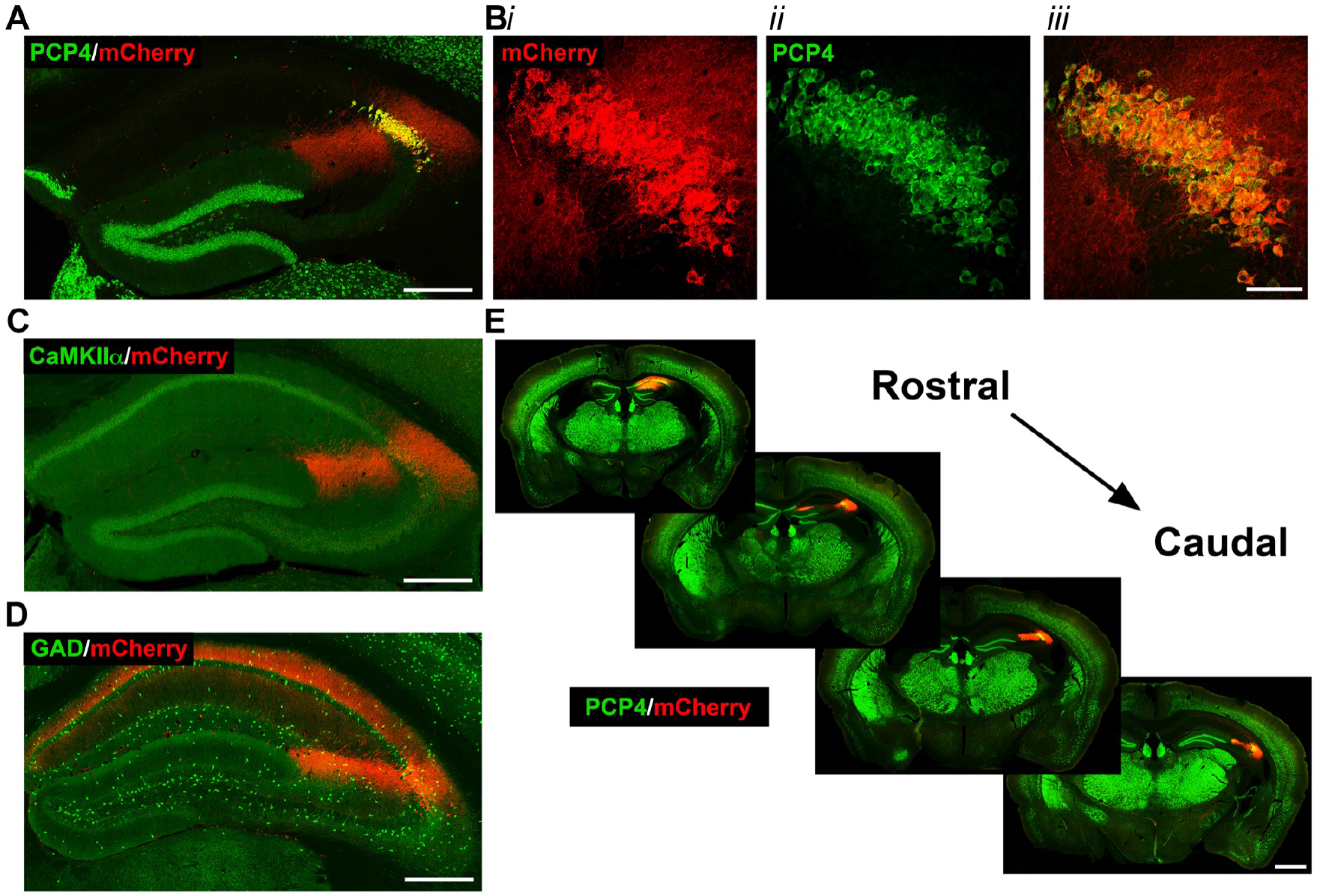
Selective expression of mCherry-tagged DREADD receptors in CA2 pyramidal cells of *Amigo2*-icreERT2 mice. Coronal sections from *Amigo2-icreERT2+* mice infused unilaterally with AAV-hSyn-DIO-hM3D(Gq)-mCherry (hM3Dq AAV; A-C,E) or bilaterally with AAV-hSyn-DIO-hM4D(Gi)-mCherry (hM4Di AAV; D) and treated with tamoxifen. A-B. Expression of hM3Dq-mCherry and the CA2-specific marker PCP4, in the hippocampus (A) and CA2 (B). No evidence of mCherry expression was found outside of the PCP4 expression region, either toward CA1 or CA3. In B, *i* shows DREADD-mCherry expression, *ii* shows PCP4 expression and *iii* shows the merged image. Expression of DREADD-mCherry colocalizes with CaMKIIα, a marker for principal neurons in hippocampus (C), but does not colocalize with GAD, a marker for inhibitory neurons (D). Note that hM4Di-mCherry (shown in D) fills axons projecting to CA1. (E) Expression of hM3Dq-mCherry colocalizes with expression of PCP4 across the rostral to caudal extent of CA2. Scale bars = 200 μm (A, C, D), 50 μm (B) and 1 mm (E).

With genetic access to CA2 pyramidal cells gained, we could selectively modify activity of CA2 neurons *in vivo* with excitatory or inhibitory DREADDs and measure the resulting network and behavioral effects. One advantage of DREADDs is that compared with tetanus toxin light chain, which permanently silences neuronal output, DREADDs permit transient modification of neuronal activity, reducing the potential for compensatory circuit reorganization.

To measure the effect of increasing CA2 neuronal activity on hippocampal and prefrontal cortical population oscillatory activity, *Amigo2*-icreERT2+ and control *Amigo2*-icreERT2-mice were infused unilaterally with hM3Dq AAV, treated with tamoxifen and then implanted with electrodes in hippocampus and PFC (see Fig. S4). To confirm that hM3Dq increased neuronal activity, single-unit firing rate was measured from CA2/proximal CA1 pyramidal neurons. CNO treatment dose-dependently increased the firing rate of pyramidal neurons following CNO administration (Fig. S5). Next, *Amigo2*-icreERT2+ and control *Amigo2*-icreERT2-mice were treated with various doses of CNO or vehicle as control, and hippocampal LFPs were assessed for CNO treatment-dependent effects using spectral analyses, focusing on theta (5-10 Hz), beta (14-18 Hz) low gamma (30–60 Hz) and high gamma (65-100 Hz) oscillations. We measured oscillatory power during the 30 to 60 min time window following treatment during each of running and resting behavioral periods (Fig. 2). We found a significant increase in low gamma power following CNO administration during running for all doses tested (N=8; F(1.904, 13.33)=9.457, *p*=0.0030, repeated-measures one-way ANOVA with Geisser-Greenhouse correction for unequal variance; 0.5 mg/kg: *p*=0.0286; 1 mg/kg: *p*=0.0286; 2 mg/kg: *p*=0.0286; 4 mg/kg: *p*=0.0191, Holm-Sidak *post hoc* test for multiple comparisons versus vehicle; Fig 2Biv). We also measured theta phase, low gamma amplitude coupling via modulation index from hippocampal recordings during periods of running, but we found no significant change in the modulation index across treatments (N=8, F(2.312, 16.19)=2.376, *p*=0.1188, repeated-measures one-way ANOVA with Geisser-Greenhouse correction for unequal variance; data not shown). During periods of rest, we also found a significant increase in low gamma power following CNO administration (F(2.306, 16.15)=32.2, *p*<0.0001, repeated-measures one-way ANOVA with Geisser-Greenhouse correction for unequal variance; 0.5 mg/kg: *p*=0.1008; 1 mg/kg: *p*=0.0161; 2 mg/kg: *p*=0.0002; 4 mg/kg: *p*=0.0004, Holm-Sidak *post hoc* test for multiple comparisons versus vehicle; Fig 2Civ). During periods of rest, beta power was significantly decreased following CNO treatment compared with vehicle (F(1.408, 9.857)=10.07, *p*=0.0066; repeated-measures one-way ANOVA with Geisser-Greenhouse correction for unequal variance; 0. 5 mg/kg: *p*=0.0486; 1 mg/kg: *p*=0.1240; 2 mg/kg: *p*=0.0545; 4 mg/kg: *p*=0.0133, Holm-Sidak *post hoc* test for multiple comparisons versus vehicle; Fig 2C*iii*). High gamma power was not significantly changed by CNO treatment compared with vehicle during either run (F(1.384, 9.69)=2.288, *p*=0.1602, repeated-measures one-way ANOVA with Geisser-Greenhouse correction for unequal variance, Fig 2Bv) or rest (F(1.286, 9.003)=4.775, *p*<0.0501, repeated-measures one-way ANOVA with Geisser-Greenhouse correction for unequal variance, Fig. 2Cv). In contrast, in control *Amigo2*-icreERT2-mice, during periods of running, CNO treatment had no effect on low or high gamma power (N=4; low gamma: F(1.669, 5.006)=1.36, *p*=0.3281; high gamma: F(1.895, 5.684)=0.5079, *p*=0.6175, repeated-measures one-way ANOVA with Geisser-Greenhouse correction for unequal variance; Fig 2D, Fig. S6).

**Figure 2.**
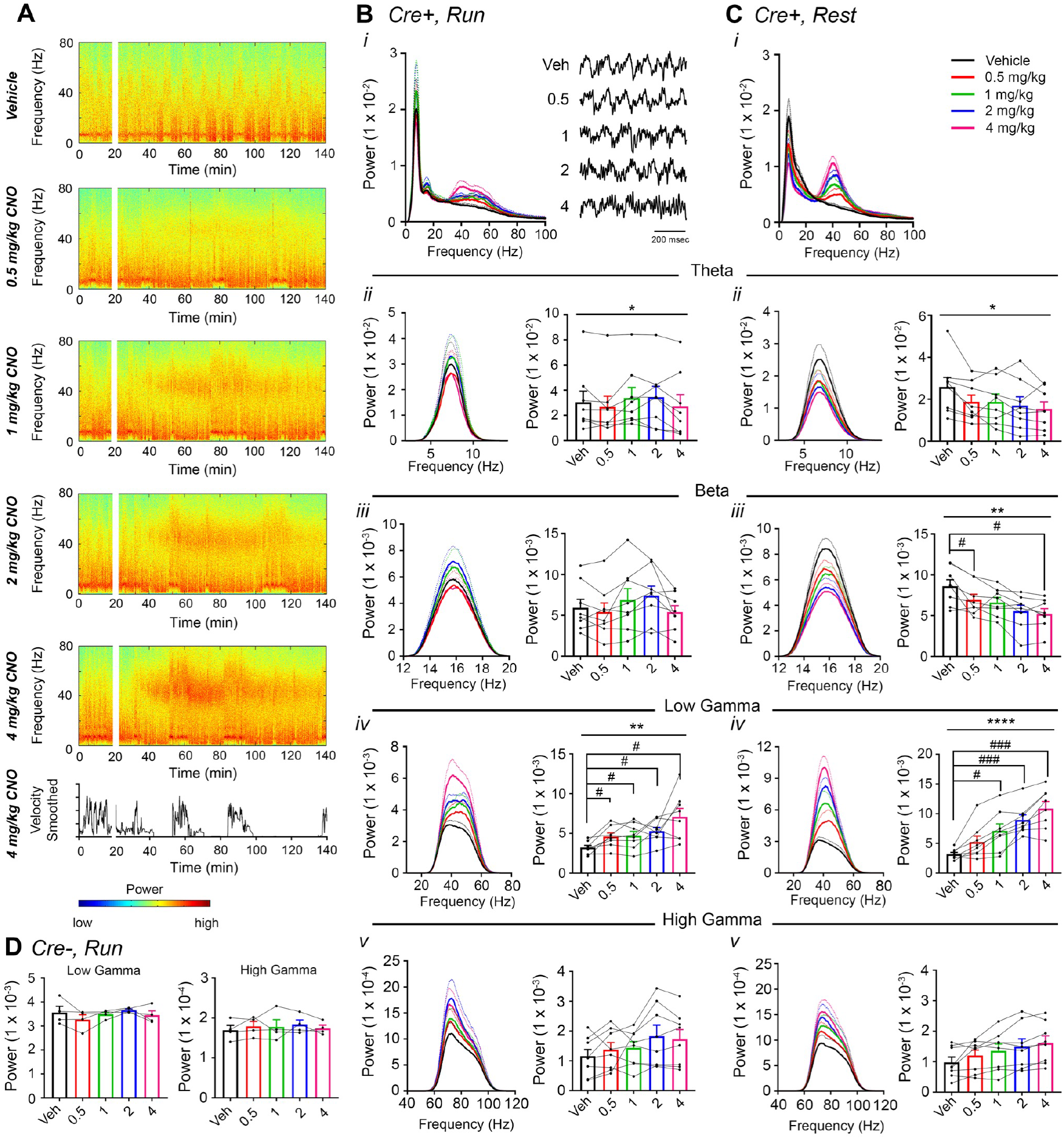
CNO treatment dose-dependently increases low gamma power in hippocampus of hM3Dq-infused *Amigo2-icreERT2+* mice. (A) Spectrograms of hippocampal LFP recordings depicting LFP power according to different frequencies over time. Vehicle/CNO administration time is shown by the white bar, and the treatment is shown to the left of each spectrogram. Locomotor velocity is shown in the bottom panel, corresponding to the 4 mg/kg CNO spectrogram. (B-C) Power measures for hippocampal LFPs in *Amigo2-icreERT2+* mice during periods of running (B) and resting (C). For each of B and C: (i) Power spectral density plots from raw LFPs for frequencies up to 100 Hz. *(ii-v)* Power spectral density plots and peak power from LFPs filtered in the theta (5-10 Hz; *ii)*, beta (14-18 Hz; *iii)*, low gamma (30-60 Hz; *iv)*, or high gamma (65100 Hz; v) frequency ranges. In Bi, raw LFP traces are shown to the right of the power spectral density plot for each treatment, sampled during a period of running during the 30-60 minutes following treatment listed. In *Bii-v* and *Cii-v*, plots on the left show power spectral density plots for each frequency band, and plots on the right show mean peak power for the population of animals in colored bars and data from individual animals as black dots. (Bii) Theta power varied significantly upon treatment during running (N=8 mice (3 female, 5 male); Friedman statistic=11.3; *p*=0.0234, results of *post hoc* tests not significant). *B(iii)* CNO treatment did not significantly affect beta power (F(2.274, 15.91)=2.91, *p*=0.0784, repeated-measures one-way ANOVA with Geisser-Greenhouse correction for unequal variance). (B*iv*) CNO treatment produced a significant dose-dependent increase in low gamma power during running (F(1.904, 13.33)=9.457, *p*=0.0030, repeated-measures one-way ANOVA with Geisser-Greenhouse correction for unequal variance; results of Holm-Sidak *post hoc* tests are shown by symbols. (Bv) CNO treatment did not significantly affect high gamma power during running (F(1.384, 9.69)=2.288, *p*=0.1602, repeated-measures one-way ANOVA with Geisser-Greenhouse correction for unequal variance). (C*ii*) Theta power varied significantly upon treatment during rest (same N; F(1.972, 13.81)=4.825, *p*=0.0261, repeated-measures one-way ANOVA with Geisser-Greenhouse correction for unequal variance; results of *post hoc* tests not significant). *(Ciii)* CNO treatment produced a significant decrease in beta power during rest (F(1.408, 9.857)=10.07, *p*=0.0066, repeated-measures oneway ANOVA with Geisser-Greenhouse correction for unequal variance; results of Holm-Sidak *post hoc* tests are shown by symbols. (C*iv*) CNO treatment produced a significant dose-dependent increase in low gamma power during rest (F(2.306, 16.15)=32.2), p<0.0001, repeated-measures one-way ANOVA with Geisser-Greenhouse correction for unequal variance; results of Holm-Sidak *post hoc* tests are shown by symbols). (C*v*) CNO treatment did not significantly affect high gamma power during rest (F(1.286, 9.003)=4.775, *p*=0.0501, repeated-measures one-way ANOVA with Geisser-Greenhouse correction for unequal variance). (D) Peak low gamma (left plot) and high gamma (right plot) power for the population of *Amigo2*-icreERT2-mice infused with hM3Dq, treated with tamoxifen and challenged with CNO. Neither low gamma power nor high gamma power changed significantly in response to CNO administration during running (N=4 male mice; low gamma: F(1.669, 5.006)=1.36, *p*=0.3281; high gamma: F(1.895, 5.684)=0.5079, *p*=0.6175, repeated-measures one-way ANOVA with Geisser-Greenhouse correction for unequal variance). \**p*<0.05, \*\**p*<0.01, \*\*\*\**p*<0.0001, one-way ANOVA; #*p*<0.05, ###*p*<0.001, Holm-Sidak *post hoc* test.

Given the role of the hippocampal-prefrontal cortical pathway in spatial working memory and the involvement of gamma synchrony between the two structures^30^ as well as the previous finding that gamma synchrony is impaired in a mouse model of schizophrenia^25^, we wondered what contribution CA2 activity makes toward PFC gamma oscillations. Therefore, we asked whether hippocampal low gamma oscillations resulting from CA2 activation could be detected in PFC (Fig. 3). Using dual recordings from hippocampus and PFC, with implanted wire electrodes targeting prelimbic cortex (see Fig. S4), we found that CNO treatment induced significant increases in low gamma power in PFC during both run and rest periods (N=4; run: F(1.168, 3.505)=9.149, *p*=0.0450; rest: (1.561, 4.684)=4.684, *p*=0.0409, repeated-measures one-way ANOVA with Geisser-Greenhouse correction for unequal variance, Fig. 3B-C). Theta, beta and high gamma powers were not affected by CNO treatment (data not shown). Control *Amigo2*-icreERT2-animals showed no significant change in PFC low gamma power following CNO administration (N=4; run: F(1.349, 4.047)=1.809, *p*=0.2617; Fig. 3D and Fig. S7). Further, we detected no significant changes in low gamma power in *Amigo2*-icreERT2+ animals implanted with wire electrodes that missed their PFC target (N=3; run: F(1.742, 3.483) =0.7609, *p*=0.5145; repeated-measures one-way ANOVA with Geisser-Greenhouse correction for unequal variance; Fig. 3E, Fig. S4C; Fig. S8) despite those animals showing increased low gamma power in hippocampus (N=3; f(1.39, 2.781)=81.51, *p*=0.0036; repeated-measures one-way ANOVA with Geisser-Greenhouse correction for unequal variance). These findings indicate that the increase in gamma power we detected in PFC was not due to electrical artifact or brain-wide changes in activity but rather to specific hippocampal inputs into the PFC^31,32^.

**Figure 3.**
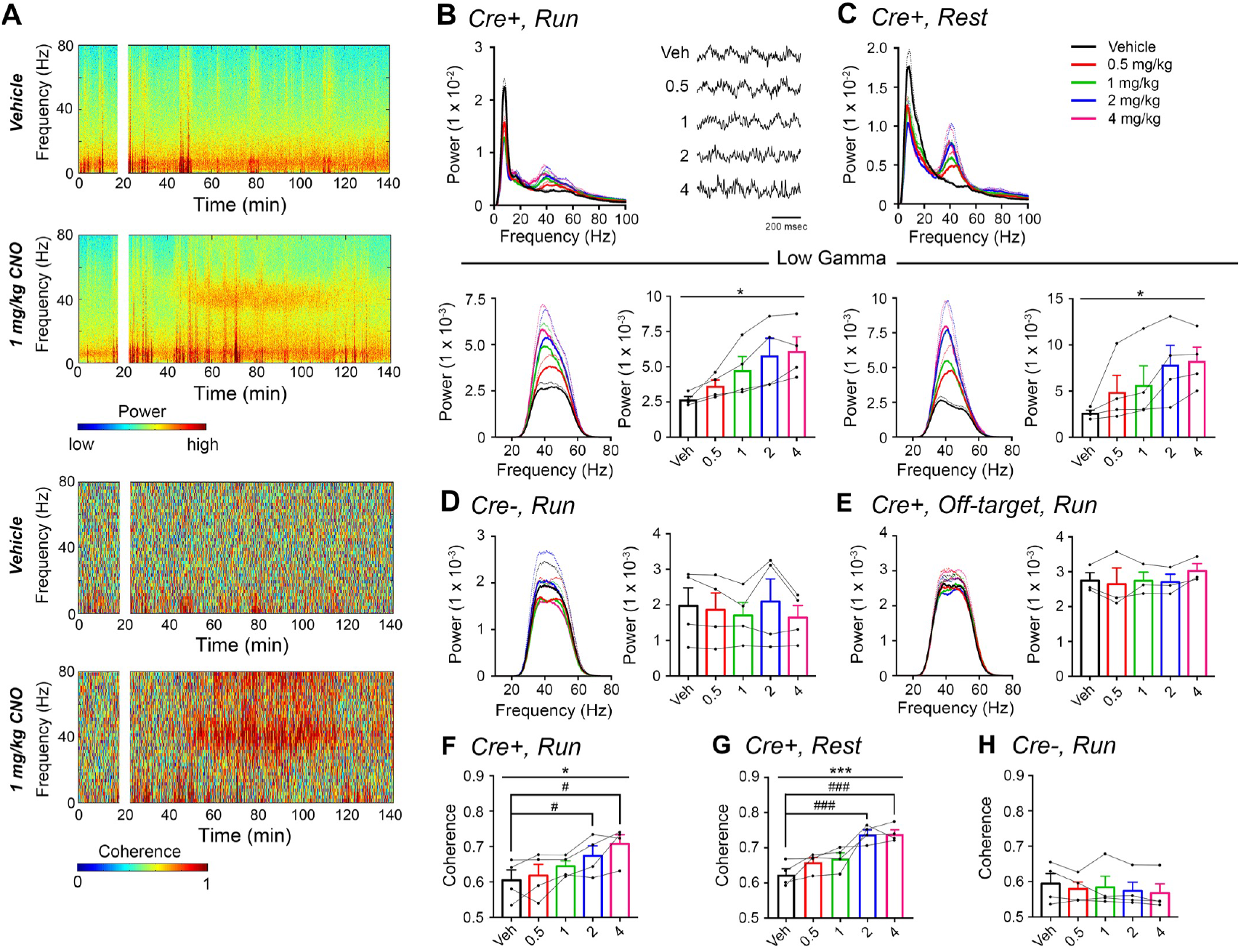
CNO treatment dose-dependently increases low gamma power in PFC of hM3Dq-infused *Amigo2-icreERT2+* mice. (A) Spectrograms of PFC LFP recordings depicting power (top two panels) and coherograms depicting coherence between PFC and hippocampal LFP recordings (bottom two panels) according to different frequencies over time. Vehicle/CNO (1 mg/kg, SQ) administration time is shown by the white bar, and the treatment is shown to the left of each spectrogram. (B-C) Power measures for PFC LFPs during periods of running (B) and resting (C) for *Amigo2-icreERT2+* mice. For each of B and C: Top plots show power spectral densities of LFPs for frequencies up to 100 Hz and bottom plots show power measured from PFC LFPs filtered in the low gamma (30-60 Hz) frequency range for each of run and rest periods. LFP traces in B show example LFPs during periods of running following the listed treatment. CNO treatment significantly increased low gamma power during both running (N=4 mice (3 male, 1 female); F(1.168, 3.505)=9.146), *p*=0.0450, repeated-measures one-way ANOVA with Geisser-Greenhouse correction for unequal variance; results of Holm-Sidak *post hoc* tests not significant) and resting (F(1.561, 4.684)=7.155), *p*=0.0409, repeated-measures one-way ANOVA with Geisser-Greenhouse correction for unequal variance). (D) Mean gamma power spectra and peak gamma power recorded from PFC for the population of hM3Dq infused *Amigo2*-icreERT2-mice during periods of running. Low gamma power did not significantly change in *Amigo2*-icreERT2-mice upon CNO administration (N=4 male mice; F(1.349, 4.047)=1.809, *p*=0.2617; repeated-measures one-way ANOVA with Geisser-Greenhouse correction for unequal variance). (E) Mean low gamma power spectra and peak low gamma power for recordings from *Amigo2-icreERT2+* mice infused with hM3Dq in which recording wires missed the target PFC area. CNO administration produced no significant change in peak low gamma power from off-target recordings (N=3 mice (1 male, 2 female); F(1.742, 3.483) =0.7609, *p*=0.5145; repeated-measures one-way ANOVA with Geisser-Greenhouse correction for unequal variance). Each of the animals used for data shown in E showed increased low gamma power in hippocampus upon CNO administration. (F-G) Mean coherence between PFC and hippocampal low gamma-filtered LFPs during periods of run (F) and rest (G) for *Amigo2-icreERT2+* mice successfully targeted to PFC. CNO treatment produced a significant increase in low gamma coherence between hippocampus and PFC during both running (N=4 mice; F(1.595, 4.786)=8.279, *p*=0.0305; repeated-measures one-way ANOVA with Geisser-Greenhouse correction for unequal variance, results of Holm-Sidak *post hoc* tests shown by symbols) and resting (F(4, 12)=11.71, *p*=0.0004; repeated-measures one-way ANOVA, results of Holm-Sidak *post hoc* tests shown by symbols). (H) *Amigo2*-icreERT2-animals showed no significant change in low gamma coherence upon CNO administration (N=4 male mice; F(4, 12)=1.053, *p*=0.4209; repeated-measures one-way ANOVA). All spectral plots show mean spectra for the population of animals with colors representing treatments. Bar graphs show mean peak gamma power (B-E) or mean gamma coherence (F-H) for the population of animals in colored bars according to treatment and data from individual animals in black dots. Dotted lines on spectral plots and error bars on bar graphs represent standard error of the mean. \**p*<0.05, \*\*\**p*<0.001, one-way ANOVA; #*p*<0.05, ### *p*<0.001, Holm-Sidak *post hoc* test.

Because we found increased low gamma power upon CNO administration in both hippocampus and PFC, we analyzed LFP coherence between the two signals to measure the extent to which the two brain areas oscillated together. CNO administration produced a significant increase in low gamma coherence between hippocampus and PFC during both run (N=4; f(1.595, 4.786)=8.279, *p*=0.0305; repeated-measures one-way ANOVA with Geisser-Greenhouse correction for unequal variance; 0.5 mg/kg: *p*=0.5808, 1 mg/kg: *p*=0.2079, 2 mg/kg: *p*=0.0292, 4 mg/kg: *p*=0.0292; Holm-Sidak *post hoc* tests versus vehicle; Fig. 3F), and rest (F(4, 12)=11.71, *p*=0.0004; repeated measured one-way ANOVA; 0.5 mg/kg: *p*=0.1189, 1 mg/kg: *p*=0.1018, 2 mg/kg: *p*=0.0006, 4 mg/kg: *p*=0.0006; Holm-Sidak *post hoc* tests versus vehicle; Fig. 3G). In contrast, treatment with CNO produced no significant change in coherence between hippocampus and PFC in control *Amigo2*-icreERT2-animals (N=4; F(4, 12)=1.053, *p*=0.4209; repeated-measures one-way ANOVA; Fig. 3H).

### Increasing CA2 pyramidal cell activity decreases sharp-wave ripple oscillations

CA2 neuronal activity was recently reported to ramp up before the onset of sharp-wave ripples^8^, so we were interested in whether and how modifying CA2 neuronal activity would impact sharp-wave ripples recorded in CA1. Therefore, we measured ripple oscillations from the CA1 pyramidal cell layer of *Amigo2-* icreERT2+ and control *Amigo2*-icreERT2 mice infused with hM3Dq during periods of rest 30-60 minutes following administration of either CNO (0.5 mg/kg, SQ; Fig. 4) or vehicle as control. We chose to use a low dose of CNO in this experiment to minimize the possibility that ripple-filtered LFPs would be contaminated by neuronal spiking in response to CNO administration independent of ripple-associated spiking. In *Amigo2*-icreERT2+ animals, CNO administration significantly decreased ripple event rate relative to that observed following vehicle administration (N=8; t(7)=4.574, *p*=0.0026; two-tailed paired t-test; Fig. 4C), although ripple amplitude was not significantly affected (t(7)=0.3004, *p*=0.7726; two-tailed paired t-test; Fig. 4D). In control *Amigo2*-icreERT2-animals, CNO administration had no effect on ripple event rate or amplitude (N=4; event rate: t(3)=1.871, *p*=0.1581; amplitude: t(3)=0.3193, *p*=0.7704; two-tailed paired t-test; Fig. 4E-F). Further experiments in a different line of hM3Dq-expressing animals are presented in Fig. S13-S17.

**Figure 4.**
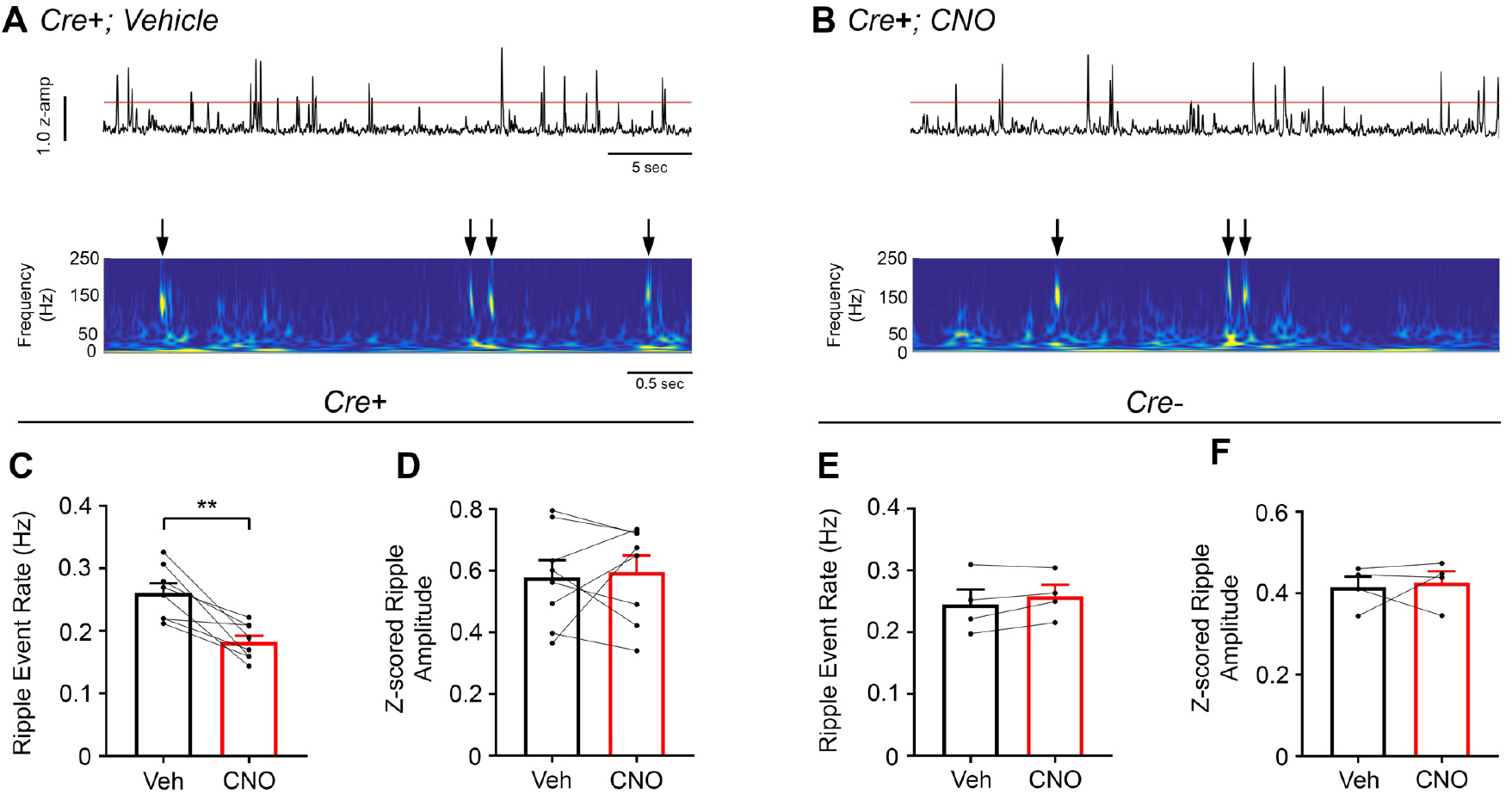
Chemoactivation of CA2 pyramidal cells with hM3Dq decreases high-frequency ripple event rate. (A) Envelopes of ripple-filtered CA1 LFPs (top) recorded during periods of rest following administration of vehicle (left) or CNO (right; 0.5 mg/kg, SQ) and wavelet-filtered spectrograms (bottom) of the same LFPs. Cooler colors represent low power and warmer colors represent high power. Arrows denote examples of ripples shown by spectrogram. (C-D) Ripple event rate (C) but not amplitude (D) was significantly decreased in hM3Dq-expressing mice following CNO administration compared to that following vehicle administration (Ripple event rate: N=8 mice (5 male, 3 female); t(7)=4.574, *p*=0.0026; two-tailed paired t-test; Amplitude: t(7)=0.3004, *p*=0.7726, two-tailed paired t-test). (E-F) Ripple event rate and amplitude were not significantly changed in *Amigo2*-icreERT2-hM3Dq-infused mice (Ripple event rate: N=4 male mice, t(3)=1.871, *p*=0.1581, two-tailed paired t-test; Amplitude: N=4 male mice, t(3)=0.3193, *p*=0.7704, two-tailed paired t-test). \*\**p*<0.01.

### CA2 pyramidal cell inhibition decreases hippocampal and prefrontal cortical low gamma power

Based on our finding that increasing activity of CA2 neurons in hM3Dq-expressing mice increased low gamma power in hippocampus and PFC, we hypothesized that inhibition of CA2 pyramidal neurons with hM4Di would decrease gamma power. As a control experiment to ascertain whether hM4Di would decrease CA2 synaptic output in our system, we infused *Amigo2-icreERT2+* mice with AAV-EF1a-DIO-hChR2(H134R)-EYFP (ChR2) and hM4Di AAVs, treated animals with tamoxifen, and then implanted the animals with fiber optic probes in CA2 and electrode bundles in the ipsilateral intermediate CA1. Optogenetic stimulation of CA2 in these awake, behaving animals evoked detectable voltage responses in CA1. Following CNO administration (5 mg/kg, SQ), the amplitude of the light-evoked responses was decreased to 5% of pre-CNO response amplitudes as early as 20 min post CNO treatment (the earliest we tested). In this preparation, we detected inhibition of CA2 responses for 4 hours. By 24 hours, responses recovered to 77.20% of pre-CNO response amplitude (Fig. S9).

To test our hypothesis that hM4Di inhibition of CA2 output would decrease hippocampal and prefrontal cortical low gamma power, we recorded LFPs from *Amigo2*-icreERT2+ and control *Amigo2*-icreERT2-mice infused with hM4Di AAV, treated with tamoxifen and implanted with electrodes. Hippocampal LFPs were measured from the primary target of CA2 pyramidal neurons, CA1 (4 mice with dorsal CA1 electrodes, 4 mice with intermediate CA1 electrodes, Fig. S10), because the majority of the neuronal inhibition by hM4Di occurs at the axon terminal to reduce neurotransmitter release^33^. Using identical analyses as for hM3Dq-infused animals, we compared LFPs filtered in the theta (5-10 Hz), beta (14-18 Hz), low gamma (30-60 Hz) and high gamma (65-100 Hz) frequency ranges during periods of running and resting 30-60 minutes following administration of CNO (5 mg/kg, SQ) or vehicle. We found a significant decrease in low gamma power during running following CNO administration compared with vehicle (t(7)=4.408, *p*=0.0031, two-tailed paired t-test, Fig. 5AiV, Fig. 5F). However, modulation index, a measure of theta phase, gamma amplitude coupling, was not significantly affected by CNO administration during running (t(7)=2.07, *p*=0.0772; twotailed paired t-test, data not shown). We also found a significant increase in beta power during running following CNO administration compared with vehicle (t(7)=2.401, *p*=0.0474, two-tailed paired t-test, Fig. 5A*iii*). Treatment with CNO did not affect theta or high gamma power during running and did not affect power in any of these frequency bands during periods of rest (Fig. 5A-B).

**Figure 5.**
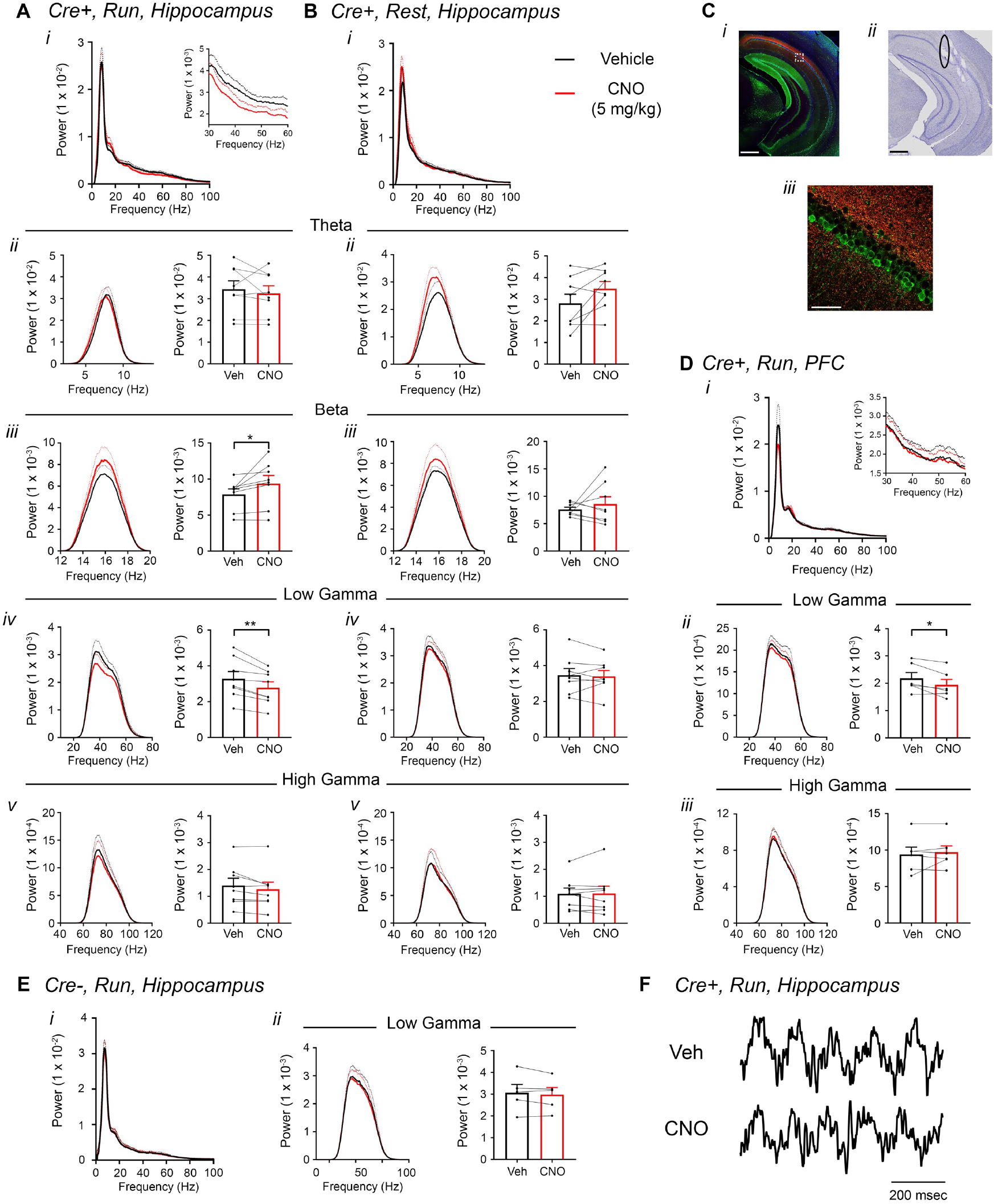
Inhibition of CA2 pyramidal cells with hM4Di decreases hippocampal and PFC low gamma power. (A-B) Hippocampal LFP power measures from *Amigo2-icreERT2+* mice infused with hM4Di AAV and treated with vehicle or CNO (5 mg/kg, SQ; LFP samples 30-60 minutes following treatment) during periods of running (A) and resting (B). For each of A and B: *(i)* Power spectral density plots from raw LFPs for frequencies up to 100 Hz. Inset plot in A*i* is expanded from A*i*. *(ii-v)* Power spectral density plots and peak power measured in the theta (5-10 Hz; *ii*), beta (14-18 Hz; *iii*), low gamma (30-60 Hz; *iv*) or high gamma (65-100 Hz; v) frequency ranges. In *Aii-v* and *Bii-v*, plots on the left show power spectral density for the listed frequency bands, and plots on the right show mean peak power for the population of animals in colored bars according to treatment and dots representing data from individual animals. (A*iii*) CNO administration produced a significant increase in beta power during running (N=8 mice (4 female, 4 male); t(7)=2.401, *p*=0.0474, two-tailed paired t-test. (A*iv*) CnO administration produced a significant decrease in hippocampal low gamma power during running (same N; t(7)=4.408, *p*=0.0031, two-tailed paired t-test). CNO treatment did not affect theta power during running (t(7)=0.7786, *p*=0.4617; A*ii*), high gamma power during running (t(7)=2.029, *p*=0.0821; Av), theta power during rest (t(7)=2.214, *p*=0.0625; B*ii*), beta power during rest (t(7)=0.7453, *p*=0.0625; B*iii*), low gamma power during rest (t(7)=0.4522, *p*=0.6648; B*iv*) or high gamma power during rest (t(7)=0.172, *p*=0.8683; Bv). (C) Expression of mCherry-tagged hM4Di in intermediate CA1 and electrode tracks at a similar position. (C*i*) Expression of mCherry-tagged hM4Di (red) and calbindin (green) in an intermediate hippocampal section. The white box shows the area that is expanded in C*iii*. Axons expressing hM4Di target intermediate CA1, with preferential targeting toward *stratum oriens*. (C*ii*) Electrode tracks of intermediate CA1 recording wires (black ellipse surrounds one of the tracks). (D) PFC LFP power measures from same mice used in A-B. (*i*) Power spectral density plots from raw LFPs for frequencies up to 100 Hz. Inset plot is expanded from the adjacent plot. (*ii-iii*) Power spectral density plot and peak power measured from low gamma (*ii*) and high gamma (*iii*) filtered LFPs. CNO administration produced a significant decrease in PFC low gamma power during running (N=6 mice (3 female, 3 male); t(5)=2.948, *p*=0.0320, two-tailed paired t-test) but did not affect PFC high gamma power (t(5)=0.738, *p*=0.4937. (E) Power spectral density plot for the population of *Amigo2*-icreERT2-mice infused with hM3Di, treated with tamoxifen and challenged with CNO. Plots show spectral density of frequencies below 100 Hz (*i*), low gamma-filtered LFP spectral power and peak low gamma power for the population of animals (*ii*). CNO administration did not significantly affect low gamma power in *Amigo2*-icreERT2-mice during running (N=5 male mice; t(4)=1.079, *p*=0.3413; two-tailed paired t-test). (F) Example LFP traces from periods of running following vehicle or CNO treatment. \**p*<0.05, \*\**p*<0.01. Scale bars = 500 um (C*i*, *ii*) and 75 um (C*iii*).

CA2 pyramidal neurons have been shown to possess axons with large rostral to caudal trajectories, primarily targeting CA1^34^. Consistent with a projection toward caudal CA1, we observed fluorescence from hM4Di-mCherry+ axon fibers in caudal intermediate CA1, with most of the fluorescently-labeled CA2 axons targeting *stratum oriens* in CA1^29^ (Fig. 5C). CA1 neurons, in turn, project to PFC^32,35,36^. Therefore, we asked whether inhibition of CA2 pyramidal neurons would impact low gamma power recorded in PFC. A subset of *Amigo2*-icreERT2+ and control *Amigo2*-icreERT2-mice with electrodes in CA1 were also implanted with electrodes in PFC and treated with CNO (5 mg/kg, SQ) or vehicle control. In *Amigo2*-icreERT2+ mice, we observed a significant decrease in PFC low gamma power during running following CNO administration compared with vehicle (N=6; t(5)=2.948, *p*=0.0320; two-tailed paired t-test; Fig. 5D), suggesting that CA2 activity modulates PFC low gamma oscillations, likely via intermediate CA1. Control *Amigo2-*icreERT2-animals showed no significant change in hippocampal or PFC low gamma power in response to CNO treatment compared with vehicle (N=5; hippocampus: t(4)=1.079, *p*=0.3413, two-tailed paired t-test; PFC: t(4)=0.4293, *p*=0.6898, two-tailed paired t-test; Fig. 5E).

### CA2 pyramidal cell inhibition increases hippocampal ripple oscillations

To assess the influence of inhibiting CA2 output on ripple oscillations, we measured ripples from the CA1 pyramidal cell layer in *Amigo2*-icreERT2+ and control *Amigo2*-icreERT2-mice during periods of rest, 30-60 minutes following administration of either CNO (5 mg/kg, SQ) or vehicle control. As predicted based on our findings in hM3Dq-infused animals, CNO administration significantly increased ripple event rate in hM4Di-infused animals (N=6; t(5)=3.809, *p*=0.0063; one-tailed paired t-test; Fig. 6C). CNO administration also increased ripple amplitude in hM4Di animals (t(5)=3.069, *p*=0.0278; two-tailed paired t-test; Fig. 6D). By contrast, in control *Amigo2*-icreERT2-mice, CNO administration did not significantly change ripple event rate or amplitude (N=6; event rate: *W*=5, *p*=0.6875, Wilcoxon signed-ranked test; amplitude: t(5)=0.5165, *p*=0.6275, two-tailed paired t-test; Fig. 6E-F). These data, together with our hM3Dq ripple findings, indicate that hippocampal ripple occurrence is negatively modulated by activity of CA2 pyramidal neurons.

**Figure 6.**
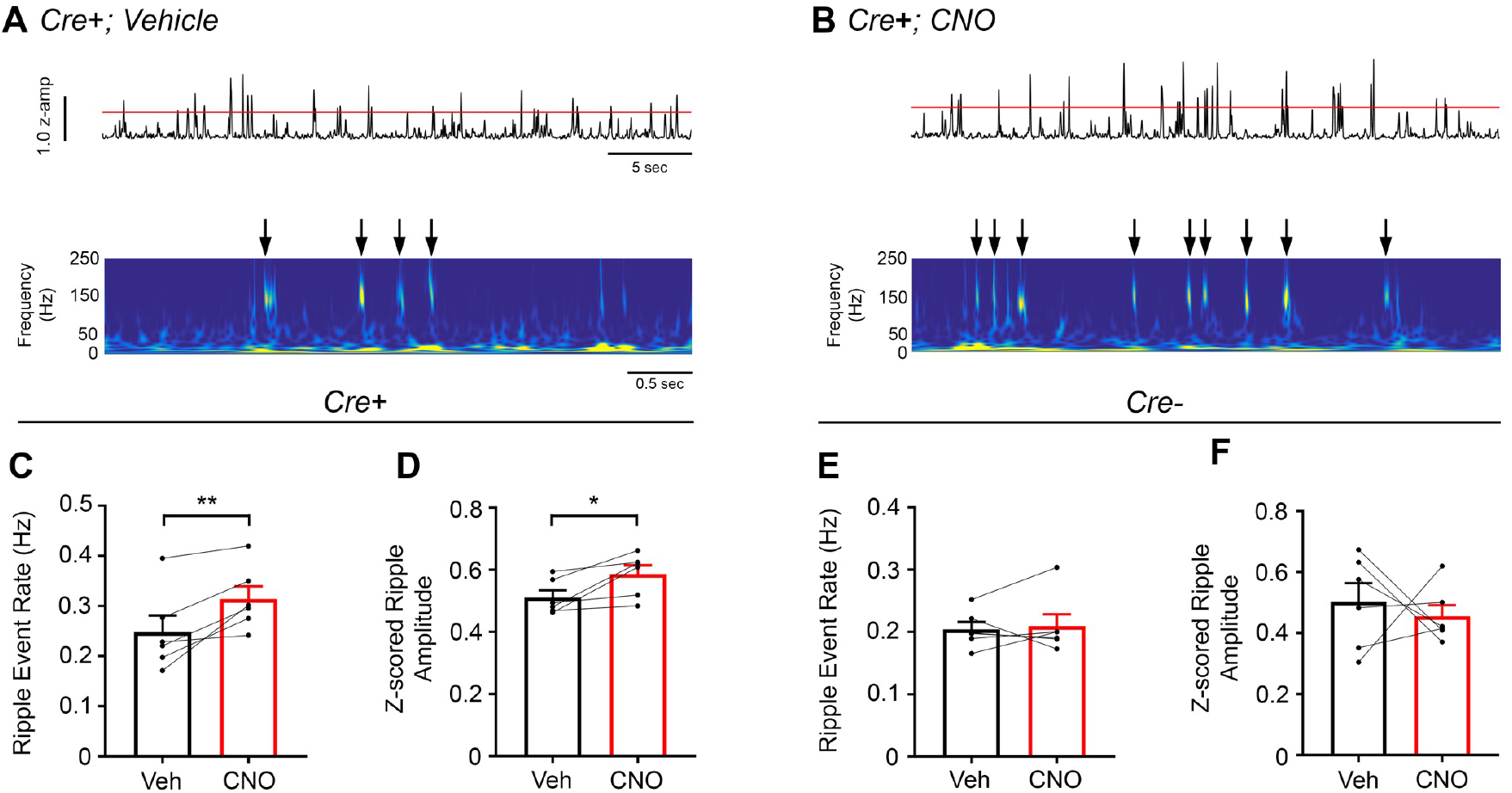
Inhibition of CA2 pyramidal cells with hM4Di increases high-frequency ripple event rate and amplitude. (A) Envelopes of ripple-filtered CA1 LFPs (top) recorded during periods of rest following administration of vehicle (left) or CNO (right; 5 mg/kg, SQ) and wavelet-filtered spectrograms (bottom) of the same LFPs. Cooler colors represent low power and warmer colors represent high power. Arrows denote examples of ripples shown by spectrogram. (C-D) Ripple event rate (C) and amplitude (D) were significantly increased in hM4Di-expressing mice following CNO treatment compared to that following vehicle treatment (Ripple event rate: N=6 mice (3 male, 3 female); t(5)=3.809, *p*=0.0063; two-tailed paired t-test; Amplitude: N=6; t(5)=3.069, *p*=0.0278, two-tailed paired t-test). (E-F) Ripple event rate and amplitude were not significantly changed in *Amigo2*-icreERT2-hM4Di-infused mice (Ripple event rate: N=6 male mice; *W*=5, *p*=0.6875, Wilcoxon signed-ranked test; Amplitude: t(5)=0.5165, *p*=0.6275, twotailed paired t-test). \**p*<0.05, \*\**p*<0.01.

### CA2 pyramidal cell inhibition decreases social preference

To determine whether oscillatory effects of inhibiting CA2 correlate with specific behavioral effects, we tested hM4Di-infused *Am*/go2-icreERT2+ and control *Amigo2*-icreERT2-mice in behavioral assays following CNO administration. Based on our findings of reduced low gamma power in hippocampus and PFC in hM4Di animals, and because previous findings report reduced low gamma coherence between hippocampus and PFC^25^ as well as increased sharp-wave ripple occurrence^37^ in animal models of schizophrenia, we focused on behaviors shown to be impaired in animal models of schizophrenia, including social behavior, prepulse inhibition and spatial working memory^38^.

Social behavior was assessed in male and female *Amigo2*-icreERT2+ and control *Amigo2*-icreERT2-mice infused with hM4Di AAV using the social approach assay following administration of CNO (5 mg/kg, IP; Fig. 7). CNO-treated control *Amigo2*-icreERT2-mice favored the social chamber over the empty chamber. However, *Amigo2*-icreERT2+ animals showed no significant preference for the social chamber upon CNO administration (male and female mice combined: main effect of chamber: F(1,22)=9.852, *p*=0.0048; main effect of genotype: F(1,22)=0.06729, *p*=0.7977, repeated measures two-way ANOVA; time spent in social versus empty chamber: Cre+: F(1, 11)=1.15; *p*=0.3057; Cre-: F(1, 11)=13.52; *p*=0.0037, within-genotype repeated measures ANOVA; Fig. 7B, Fig. S11). Further, although our experiments were not powered to detect differences due to sex, a sex-specific difference emerged: the effect of CA2 inhibition on social approach appeared to be driven exclusively by the male animals (Fig. 7C). Among females, *Amigo2*-icreERT2+ mice showed similar preference for the social chamber over the empty chamber as *Amigo2*-icreERT2-mice (N=8; main effect of chamber: F(1, 6)=106.6, *p*<0.0001; main effect of genotype: F(1, 6)=0.001545, *p*=0.9699, repeated measures two-way ANOVA; time spent in social chamber versus empty chamber: Cre+: *p*=0.0005; Cre-: *p*=0.0002, within-genotype repeated measures ANOVA). By contrast, among males, *Amigo2*-icreERT2+ mice did not show a preference for the social chamber, while *Amigo2*-icreERT2-mice did (N=16; main effect of chamber: F(1, 14)=3.123, *p*=0.0990; main effect of genotype: F(1, 14)=0.08210, *p*=0.7787, repeated measures two-way ANOVA; time spent in social chamber versus empty chamber: Cre+: *p*=0.7932; Cre-: *p*=0.0425, within-genotype repeated measures ANOVA).

**Figure 7.**
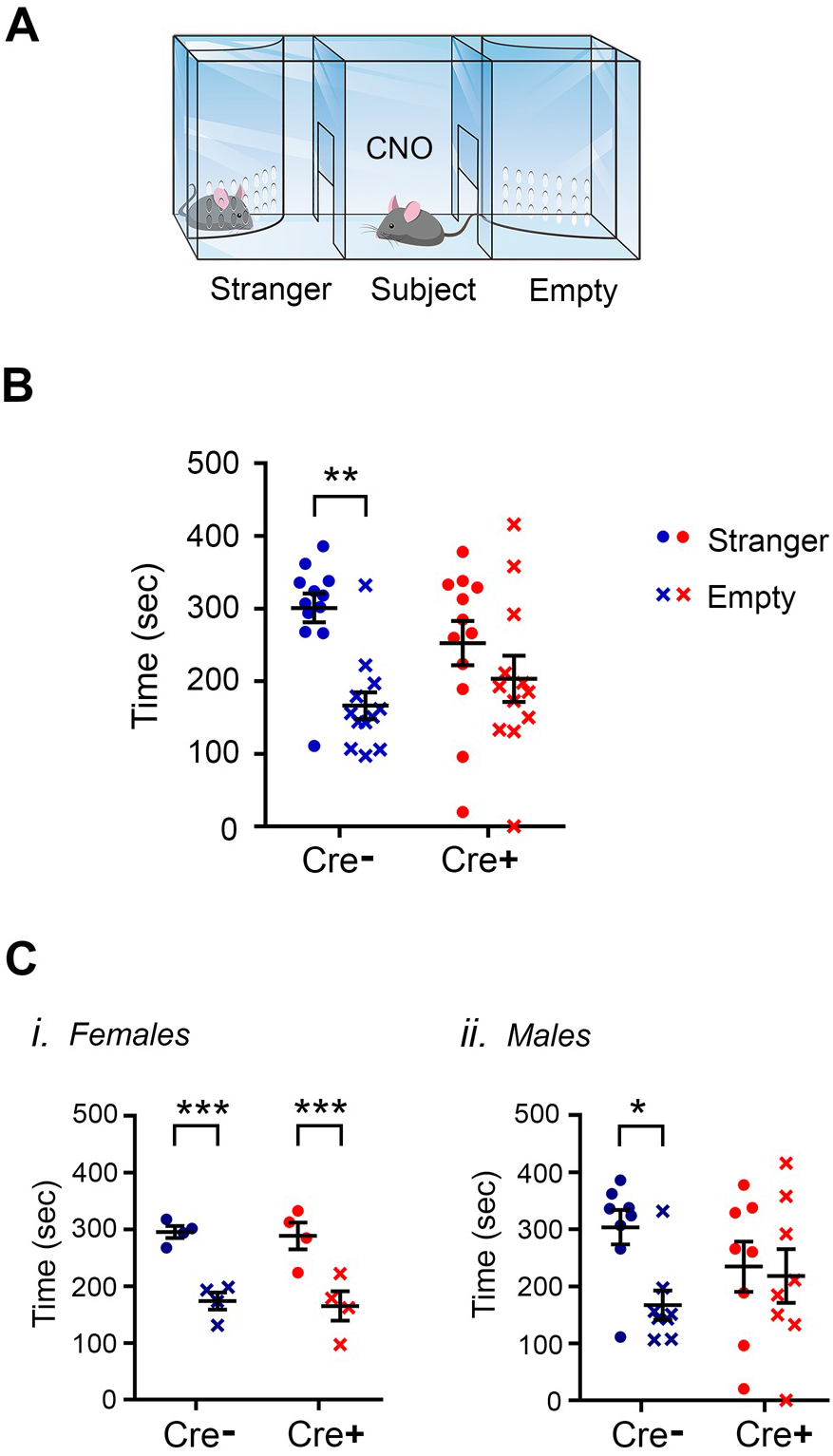
Inhibition of CA2 pyramidal cells with hM4Di decreases social approach behaviors. (A) Schematic diagram of three-chamber social approach chamber used for this study. CNO (5 mg/kg, IP) was administered to the all subject mice 20 min before starting the experiment. (B) *Amigo2*-icreERT2-, but not *Amigo2-icreERT2+*, hM4Di-expressing mice spent significantly more time in the social chamber than the empty chamber in the social approach assay (male (8 Cre+ and 8 Cre-, littermate pairs) and female (4 Cre+ and 4 Cre-, littermate pairs) mice combined; main effect of chamber: F(1, 22)=9.852, *p*=0.0048; main effect of genotype: F(1, 22)=0.06729, *p*=0.7977, repeated measures two-way ANOVA; Cre+ time spent in social chamber versus empty chamber: F(1, 11)=1.15; *p*=0.3057; Cre-time spent in social chamber versus empty chamber: F(1, 11)=13.52; *p*=0.0037, within-genotype repeated measures ANOVAs). (C) Social approach data shown for females (*i*) and males (*ii*) separately. (*i*) Female Cre+ hM4Di-expressing mice showed similar preference for the social chamber as Cre-mice (N=8 female mice (4 pairs); main effect of chamber: F(1,6)=106.6, *p*=0.0001; main effect of genotype: F(1,6)=0.001545, *p*=0.9699, repeated measures two-way ANOVA; Cre+ time spent in social chamber versus empty chamber: *p*=0.0005; Cre-time spent in social chamber versus empty chamber: *p*=0.0002, within-genotype repeated measures ANOVA). *(ii)*. Male Cre+ hM4Di-expressing mice did not show a preference for the social chamber, while Cre-mice did (N=16 male mice (8 Cre+ and 8 Cre-mice); main effect of chamber: F(1,14)=3.123, *p*=0.0990; main effect of genotype: F(1,14)=0.08210, *p*=0.7787, repeated measures two-way ANOVA; Cre+ time spent in social chamber versus empty chamber: *p*=0.7932; Cre-time spent in social chamber versus empty chamber: *p*=0.0425, within-genotype repeated measures ANOVA). Each dot represents data from an individual animal, and black horizontal lines and error bars represent means and standard errors of the mean.

Prepulse inhibition of acoustic stimuli and spatial working memory were also assessed in hM4Di-infused *Amigo2*-icreERT2+ and *Amigo2*-icreERT2-mice administered CNO (5 mg/kg, IP; Fig. S12). We failed to detect any significant differences between *Amigo2*-icreERT2+ and *Amigo2*-icreERT2-mice in any of the prepulse inhibition measures (Fig. S12A-C). To assess spatial working memory, the number and percent of spontaneous alternations were measured while animals explored a Y-maze. Again, we found no differences between *Amigo2*-icreERT2+ and *Amigo2*-icreERT2-mice in this measure of working memory (Fig. S12D-F).

## DISCUSSION

In this study, we used excitatory and inhibitory DREADDs to reversibly modify activity of CA2 pyramidal cells and examined the effect on hippocampal and prefrontal cortical oscillations and behavior. We found that increasing activity of CA2 pyramidal cells increased hippocampal and prefrontal cortical low gamma power and decreased hippocampal sharp-wave ripples. Conversely, we found that inhibiting CA2 pyramidal cell output decreased hippocampal and prefrontal cortical low gamma power and increased hippocampal sharp-wave ripples. Behaviorally, inhibiting CA2 output decreased social approach behavior. These findings demonstrate a role for hippocampal area CA2 in low gamma oscillation generation across the distributed hippocampal-prefrontal cortical network. Further, these findings support a negative regulatory role of CA2 in hippocampal sharp-wave ripples and provide further support for the role of CA2 in social behavior.

CA2 activation produced robust, dose-dependent increases in hippocampal low gamma power, and inhibition of CA2 neurons decreased low gamma power. Gamma oscillations in hippocampus reflect synchronous inhibitory postsynaptic potentials (IPSPs) from a network of interconnected perisomatically-targeted basket cells, with excitatory drive onto these interneurons arising from pyramidal cells^39–41^. The frequency of the gamma oscillations is controlled by the decay kinetics of the IPSP such that slower decay yields a lower gamma frequency^40^. Low gamma oscillations in CA1 reportedly arise from neuronal activity in CA3^10,12^ However, permanent silencing of CA3 output with tetanus toxin light chain produced only a 30% reduction in low gamma power recorded in CA1^18^, thereby challenging the notion that CA3 is the only origin of hippocampal low gamma oscillations. Here, we report that CA2 activation increases low gamma power and that acute CA2 inhibition reduces low gamma power recorded in CA1 by approximately 20%. hM4Di was previously reported to substantially, but not entirely, inhibit synaptic neurotransmitter release^33^, although we found that hM4Di produced near complete silencing of CA2 output (Fig. S9). Complete silencing of CA2 may reduce low gamma power by greater than the 20% we observed here, but more likely, silencing of both CA2 and CA3 may be needed to produce substantial inhibition of low gamma power. Based on these findings, CA2 and CA3 together likely provide the excitatory drive required to generate low gamma oscillations in CA1. Given the dynamic nature of the brain, we propose that when CA2 is inhibited, CA3, or another source, is capable of compensating, and vice versa. Further, gamma activity arising from CA2 and CA3 may engage distinct circuits involving the deep and superficial CA1 pyramidal neurons, respectively^29,42^. As such, gamma oscillations arising from the two areas may subserve distinct cognitive functions based on the output of these two populations.

Modification of CA2 neuronal activity also affected the occurrence of sharp-wave ripple oscillations. Specifically, increasing CA2 activity with hM3Dq decreased the occurrence of ripples, whereas decreasing CA2 output with hM4Di increased ripple occurrence as well as amplitude. The mechanism underlying these findings likely includes the robust inhibition that CA2 presents over CA3 neurons^9,29^. Accordingly, CA2 pyramidal cells contact local parvalbumin-expressing basket cells, which project to CA3^43^, and CA2 pyramidal cell firing is reported to discharge CA3 interneurons^8^. As a potential secondary mechanism underlying the observed inverse relationship between CA2 neuronal activity and occurrence of ripples, CA2 neurons preferentially target the deep layer of pyramidal cells in CA1^29^. During recordings of ripples from these deep CA1 pyramidal cells, dominant hyperpolarizations are observed, which contrasts with dominant depolarizations during ripples seen in superficial CA1 pyramidal cells^44^ (see also^42^). Further, stimulation of CA2 neurons produces robust feed-forward inhibitory responses onto CA1 neurons^9,44^. Therefore, silencing of CA2 neurons may remove feed-forward inhibition and produce a net excitatory response in CA1. Together, these two mechanisms may explain the significant gating influence that CA2 neurons have over hippocampal excitability and, consequently, sharp-wave ripples in CA1.

Consistent with this finding, mice in which CA2 synaptic output was fully and permanently blocked via tetanus toxin light chain showed normal ripples and also anomalous epileptiform discharges that arose from CA3 during immobility^9^. Our findings of increased ripples in CA1 upon acute inhibition of CA2 output are consistent with these findings in that in both studies, CA2 silencing increases CA3 to CA1 output during immobility. Echoing the statement by Boehringer et al^9^, our data do not agree with the suggestion by Oliva et al.^8^ that CA2 neuronal activity triggers the occurrence of ripples. Rather, CA2 activity may play a role in sculpting the CA3 network activity and gate output to CA1. Consistent with a gating, or permissive, role of CA2 toward the occurrence of ripples, Kay et al. revealed that CA2 is the only hippocampal subregion to have a substantial population of neurons that cease firing during CA1 ripples^6^. Similarly, Oliva et al. demonstrated an inverse correlation between occurrence of ripples in CA2 and CA1. During periods of low occurrence of ripples in CA2, ripple occurrence was high in CA1, and vice versa^8^. The inverse correlations described by these two findings suggest a negative regulatory role of CA2 activity on CA1 ripples, which is consistent with our findings.

Our findings also reveal a role for CA2 in beta oscillations in that CA2 activation decreased beta power whereas CA2 inhibition increased beta power. Although these oscillations have been studied far less than gamma and ripple oscillations, they are thought to contribute to hippocampal novelty detection processes; beta power is increased on exposure to a novel environment and decreases with repeated exposure^45,46^. A role for CA2 in oscillations that reflect novelty detection seems fitting given our recent findings that CA2 place fields are remapped upon exposure to novel environmental stimuli^2^ and other studies demonstrating CA2 responsiveness to novelty^5,47^.

We and others have demonstrated a significant role for CA2 in social behavior. Using mice with CA2 chronically silenced with tetanus toxin light chain, Hitti and Siegelbaum presented evidence that CA2 activity is required for one form of social recognition memory^21^. In addition, we recently demonstrated that CA2 neuron spatial representations (place fields) remap upon exposure to novel or familiar conspecific animals^2^. These effects may involve vasopressinergic signaling because deletion of one of the vasopressin receptors *(Avprlb)* that shows selective expression in CA2 results in impaired social behavior, including aggression^22^, and optogenetic stimulation of vasopressin-containing axon fibers in CA2 enhances social memory^23^. Interestingly, both vasopressin and oxytocin receptor agonists enhance Schaffer Collateral synaptic transmission in CA2^22^. Here, we report that acute inhibition of CA2 neurons with hM4Di impairs social approach behavior. Remarkably, the social approach impairment we observed may be specific to male mice in that our female, but not male, hM4Di mice behaved similar to controls. Of note, previous studies of CA2 and social behavior used only male rodents^2,21–23,48^. The mechanism underlying this dimorphic effect may include vasopressin and/or oxytocin, but whether axon fibers containing these molecules differentially innervate CA2 in males and females, or whether Schaffer Collateral synaptic potentiation^22^ or other effects induced by these peptides differ by sex is unknown. Indeed, though, sex differences in dendritic branching patterns of CA2 neurons have been reported in guinea pig^49^, so it stands to reason that other sex differences may exist in CA2.

The results of our study present further similarities between CA2 functions and impairments seen in schizophrenia. Gamma oscillations and social behavior are both impaired in patients with schizophrenia^50,51^. In addition, parvalbumin-expressing interneurons, which contribute to the generation of gamma oscillations, are notably lost from hippocampal area CA2 and PFC in tissue from patients with this disorder^52,53^. Findings from the *Df16A+/-* mouse model of schizophrenia demonstrate impaired social behavior, decreased number of parvalbumin-expressing interneurons in CA2, decreased activity of CA2 pyramidal neurons^24^, and decreased synchrony between hippocampus and PFC^25^. Additionally, the forebrain-selective calcineurin knock-out model of schizophrenia was reported to have increased CA1 ripple events during periods of resting wake^37^. We report that CA2 neuronal activity contributes to low gamma oscillations in both hippocampus and PFC, gamma coherence between hippocampus and PFC, hippocampal ripple oscillations and to social behavior, suggesting that CA2 may play a role in the pathophysiology of schizophrenia, or possibly the social deficits therein.

Here, we have provided evidence that CA2 neuronal activity bidirectionally controls hippocampal and prefrontal cortical low gamma oscillations as well as hippocampal beta and sharp-wave ripple oscillations. Further, we provide evidence that CA2 activity is required for social approach behavior, but perhaps only in male mice. These findings demonstrate a role for CA2 in the extended hippocampal-prefrontal cortical network and further support the idea that CA2 is an integral node in the hippocampal network relating to social cognition.

### Online Methods

#### Animals

Experiments were carried out in adult male and female mice (8–12 weeks at the start of experiments). Mice were housed under a 12:12 light/dark cycle with access to food and water *ad libitum*. Mice were naïve to any treatment, procedure or testing at the time of beginning the experiments described here. Mice were group-housed until the time of electrode implantation for those mice undergoing electrode implantation surgery, at which point they were singly housed. All procedures were approved by the NIEHS Animal Care and Use Committee and were in accordance with the National Institutes of Health guidelines for care and use of animals.

#### Generation of transgenic *Amigo2*-icreERT2

The BAC clone RP23-288P18 was used to generate these mice. To recombine the cDNA encoding an icreERT2 fusion protein^54^ into the BAC, we constructed a targeting vector from which we derived a targeting fragment for recombineering. The targeting fragment consisted of a 243 bp homology region (A-Box) immediately upstream of the ATG in the *Amigo2* gene. The icreERT2 cassette was fused to the A-Box replacing the *Amigo2* ATG with the icre ATG preceded with a perfect KOZAK sequence. At the 3’ end of the icreERT2 cassette a synthetic bovine growth hormone (BGH) polyadenylation signal was added after the STOP codon. For selection of recombined BACs, a flipase-site flanked neomycin resistance gene was incorporated into the targeting fragment following the icreERT2 cassette. Finally, the 3’ end of the targeting fragment contained a 263 bp homology region (B-Box) starting downstream of the *Amigo2* ATG. Recombineering was performed according to a previously described protocol^55^. In brief, the targeting fragment was electroporated into induced EL250 bacteria harboring the *Amigo2* BAC. Recombined colonies were selected on Chloramphenicol/Kanamycin plates and screened by colony PCR. The neo gene was removed from the recombined BAC by arabinose driven flipase expression.

Recombined BACs without the neo marker were linearized by restriction enzyme digestion, gel purified and electro-eluted from the gel slice. After filter dialysis with a Millipore VSWP02500 filter, the BAC fragment concentration was adjusted to 1 ng/μl and microinjected into pronuclei of B6SJLF1 mouse oocytes (Taconic, North America). Six independent founder mice resulted, which were bred to ROSA-tdTomato indicator mice. Resulting offspring that genotyped positive for both Cre and tdTomato were treated with tamoxifen (Sigma, 100 mg/kg daily administration, IP, 7 days of treatment). At least one week following the final treatment with tamoxifen, mice were perfused with 4% paraformaldehyde and brains were sectioned and examined for tdTomato expression. Two lines showed adult expression of icreERT2 in CA2; one showed sparse expression in dentate gyrus and was not used in this study, another (line 1; *B6(SJL)-Tg(Amigo2-icre/ERT2)1Ehs)* showed selective expression in CA2 within hippocampus as well as expression in fasciola cinerea and hypothalamus, among other locations (Supplementary Figure 1). Line 1 mice were used for electrophysiology, anatomy and behavioral studies here and were bred to ROSA-tdTomato (described above), GAD-eGFP, or GAD-eGFP; ROSA-tdTomato mice for histological analysis. *Amigo2*-icreERT2 mice used in this study were backcrossed to C57Bl/6 7 generations.

Genotyping of *Amigo2*-icreERT2 BAC transgenic mice was done using the following primers: BGH-F (forward primer) 5’-CTT CTG AGG CGG AAA GAA CC-3’ and dAmigo4 (reverse primer) 5’-AACTGCCCGTGGAGATGCTGG-3’. PCR protocol is 30 cycles of 94°C 30 sec., 60°C 30 sec., 72°C 30sec. PCR product is 600bp.

#### Animal Numbers

For all experiments presented, 89 *Amigo2*-icreERT2 mice (8 for histology, 50 for behavior, 29 for electrophysiology, 2 for optogenetics with electrophysiology), 13 *Amigo2*-icreERT2; ROSA-tdTomato mice (all for histology), 3 *Amigo2*-icreERT2; GAD-eGFP; ROSA-tdTomato mice (all for histology) and 4 *Amigo2*-icreERT2; GAD-eGFP mice (all for histology) were used. No statistical tests were used to determine sample sizes *a priori*, but sample sizes for histological, electrophysiological and behavioral studies were similar to those used in the field. For electrophysiology studies, *Amigo2*-icreERT2+ and *Amigo2*-icreERT2-animals were randomly selected from litters. For behavioral studies, pairs of *Amigo2*-icreERT2+ and *Amigo2*-icreERT2-animals were randomly selected from individual litters and infused with DREADD AAVs. For randomization, animals were housed with same-sex littermates following weaning but before genotyping. Genotype information was unknown at the time of randomly selecting a mouse from the cage for AAV infusion.

#### Virus infusion and tamoxifen treatment

Viruses were obtained from the viral vector core at the University of North Carolina-Chapel Hill. Mice were infused with AAV-hSyn-DIO-hM3D(Gq)-mCherry (Serotype 5; hM3Dq AAV), AAV-hSyn-DIO-hM4D(Gi)-mCherry (Serotype 5; hM4Di AAV) or equal parts AAV-EF1a-DIO-hChR2(H134R)-EYFP (Serotype 5; ChR2 AAV) and hM4Di mixed in a centrifuge tube. For virus-infusion surgery, mice were anesthetized with ketamine (100 mg/kg, IP) and xylazine (7 mg/kg, IP), then placed in a stereotaxic apparatus. An incision was made in the scalp, a hole was drilled over each target region for AAV infusion, and a 27-ga cannula connected to a Hamilton syringe by a length of tube was lowered into hippocampus (in mm: -2.3 AP, +/-2.5 ML, -1.9 mm DV from bregma). *Amigo2-* icreERT2 mice were infused unilaterally on the left side for hM3Dq AAV infusion, bilaterally for hM4Di AAV, or unilaterally on the left side for ChR2/hM4Di infusion. For each infusion, 0.5 μl was infused at a rate of 0.1 μl/min. Following infusion, the cannula was left in place for an additional 10 minutes before removing. The scalp was then sutured and the animals administered buprenorphine (0.1 mg/kg, SQ) for pain and returned to their cage. Two weeks following AAV infusion surgery, *Amigo2*-icreERT2 mice began daily tamoxifen treatments (100 mg/kg tamoxifen dissolved in warmed corn oil, IP) for a total of 7 days. At least one week following the last dose of tamoxifen, animals were euthanized and perfused with 4% paraformaldehyde for anatomical studies, or underwent electrode (and fiber optic probe for ChR2/hM4Di mice) implantation surgery, or were transferred to the University of North Carolina Mouse Behavioral Phenotyping Laboratory in Chapel Hill, NC. At least three weeks was allowed to elapse between the last dose of tamoxifen and the beginning of the behavioral studies.

#### Electrode Implantation

At least one week after the last tamoxifen treatment, mice for *in vivo* electrophysiology were implanted with electrode arrays. Mice were anesthetized with ketamine (100 mg/kg, IP) and xylazine (7 mg/kg, IP), then placed in a stereotaxic apparatus. An incision was made in the scalp, and the skull was cleaned and dried. One ground screw (positioned approximately 4 mm posterior and 2 mm lateral to Bregma over the right hemisphere) and four anchors were secured to the skull and electrode arrays were then lowered into drilled holes over the target brain regions. Electrode wires were connected to a printed circuit board (San Francisco Circuits, San Mateo, CA), which was connected to a miniature connector (Omnetics Connector Corporation, Minneapolis, MN). For all but one mouse that was implanted with tetrodes, electrodes consisted of stainless steel wire (44-μm) with polyimide coating (Sandvik Group, Stockholm, Sweden). Wires were bundled into groups of 8 and lowered to target regions. In 11 *Amigo2*-icreERT2 mice infused with hM3Dq AAV (7 Cre+, 4 Cre-) and 8 *Amigo2*-icreERT2 mice infused to hM4Di AAV (2 Cre+, 6 Cre-), electrode arrays were implanted into the left dorsal hippocampus, targeting CA2/proximal CA1 (in mm: -2.06 AP, -2.5 ML, -1.9 DV from bregma), the right dorsal hippocampus targeting CA1 (-1.94 AP, +1.5 ML, -1.5 DV from bregma), and the left PFC (+1.78 AP, -0.25 ML, -2.35 DV from bregma). In 4 *Amigo2*-icreERT2 mice infused with hM4Di AAV (4 Cre+), electrodes were lowered into left dorsal hippocampus targeting CA2/proximal CA1 (-2.06 AP, -2.5 ML, -1.9 DV from bregma), left PFC (+1.78 AP, -0.25 ML, -2.35 DV from bregma) and left intermediate hippocampus targeting CA1 (-2.92 AP, 2.75 ML, 2.1 DV from bregma). In 3 *Amigo2*-icreERT2 mice infused with hM3Dq AAV (3 Cre+), electrodes were implanted in left dorsal hippocampus targeting CA2/proximal CA1 only (-2.06 AP, -2.5 ML, -1.9 DV from bregma). In 2 *Amigo2*-icreERT2 mice infused with hM4Di AAV (2 Cre+), electrodes were implanted in left hippocampus targeting CA1 only (-1.94 AP, +1.5 ML, -1.25 DV from bregma. In one *Amigo2*-icreERT2+ infused with hM3Dq AAV, a bundle of 8 tetrodes was lowered into the left hippocampus targeting CA2/proximal CA1 (-2.06 AP, -2.5 ML, -1.9 DV from bregma) for monitoring changes in single unit firing rate upon CNO administration. In 2 *Amigo2*-icreERT2+ mice infused with ChR2/hM4Di AAV, a fiber optic probe was implanted into left CA2 (-1.95 AP, -2.25 ML, -1.65 DV from bregma) and a wire bundle was implanted into left intermediate CA1 (-3.08 AP, -2.75 ML, -2.0 DV from bregma).

#### Histology

Animals used for histology were euthanized with Fatal Plus (sodium pentobarbital, 50 mg/mL; >100 mg/kg) and underwent transcardial perfusion with 4% paraformaldehyde. Brains were then cryoprotected in 30% sucrose PBS for at least 72 hours and sectioned with a cryostat or vibratome at 40 μm.

For immunohistochemistry, brain sections were rinsed in PBS, boiled in deionized water for 3 min, and blocked for at least 1 h in 3-5% normal goat serum/0.01% Tween 20 PBS. Sections were incubated in the following primary antibodies, which have previously been validated in mouse brain^21,29^: rabbit anti-PCP4 (SCBT, sc-74186, 1:500), rabbit anti-CaMKII alpha (Abcam, ab131468, 1:250), rat anti-mCherry (Life Technologies, M11217, 1:500-1:1000), mouse anti-cre (Millipore, 3120, 1:5000), mouse anti-calbindin (Swant, D-28k, 1:500). Antibodies were diluted in blocking solution and sections were incubated for 24 h. After several rinses in PBS/Tween, sections were incubated in secondary antibodies (Alexa goat anti-mouse 488 and Alexa goat anti-rabbit 568, Alexa Goat anti-rat 568, Invitrogen, 1:500) for 2h. Finally, sections were washed in PBS/Tween and mounted under ProLong Gold Antifade fluorescence media with DAPI (Invitrogen). Images of whole-brain sections were acquired with a slide scanner using the Aperio Scanscope FL Scanner, (Leica Biosystems Inc.). The slide scanner uses a monochrome TDI line-scan camera, with a PC-controlled mercury light source to capture high resolution, seamless digital fluorescent images. Images of hippocampi were acquired on a Zeiss 780 meta confocal microscope using a 40× oil-immersion lens. Counts were made of cells expressing the Cre-dependent tdTomato fluorescent reporter. Five *Amigo2-icreERT2+;* ROSA-tdTomato +/-mice were used for this analysis with 3-5 50-μm sections per animal spanning the anterior-posterior extent of CA2. Sections were stained for PCP4 and colocalization of PCP4 with tdTomato was assessed in a total of 5,248 cells.

#### Neurophysiological data acquisition and behavioral tracking

Neural activity was transmitted via a 32-channel wireless 10× gain headstage (Triangle BioSystems International, Durham, NC) and was acquired using the Cerebus acquisition system (Blackrock Microsystems, Salt Lake City, UT). Continuous LFP data were band-pass filtered at 0.3–500 Hz and stored at 1, 000 Hz. Single unit data were sampled at 30 kHz and high-pass filtered at 250 Hz. Neurophysiological recordings were referenced to a silver wire connected to a ground screw secured in the posterior parietal bone (approximately 4 mm posterior to bregma, 2 mm lateral). To confirm that gamma power activity recorded in hippocampus and PFC were not artifacts of differential recording between the active electrode and the ground screw, in some animals, one wire per bundle targeting hippocampus or PFC, was positioned either in the cortex above hippocampus or in the striatum lateral to PFC. Referencing signals to these short or lateral wires showed LFPs that increased or decreased in gamma power upon CNO administration to hM3Dq or hM4Di-infused mice, respectively, similar to recordings that were referenced to the ground screw. For behavioral tracking, the X and Y coordinates in space of a light-emitting diode (for use with color camera) or a small piece of reflective tape (for use with infrared camera) present on the wireless headstage were sampled at 30 Hz using Neuromotive Software (Blackrock Microsystems) and position data were stored with the neural data.

For recordings from mice infused with hM3Dq AAV, baseline data was acquired for at least 20 minutes followed by treatment with vehicle (10% DMSO in saline) or CNO (0.05-4 mg/kg CNO dissolved in DMSO to 50 mM then suspended in saline, SQ) and recording continued for an additional 2 hours. During the entire recording time, mice were inside of an open field arena, which was a custom-built, 5-sided (open top) dark arena (approximately 80 cm long x 80 cm wide x 100 cm high). The walls and floor of the arena were constructed from black-colored Plexiglass. Mice administered various doses of CNO were first treated with vehicle and then increasing doses of CNO at three-day intervals. Room light remained illuminated, but a curtain was placed around the open field chamber during recordings. For neurophysiology experiments on hM4Di AAV-infused mice, after connecting headstages to the animals’ electrodes, animals were administered either vehicle or CNO (5 mg/kg, SQ) then returned to their cage for 30 minutes before starting recording. Room lights were turned off and red lights were illuminated after administering vehicle or CNO. After 30 minutes had elapsed, mice were placed in the open field arena for recording. Gamma power measurements were made during periods when the animals were moving at ≥ 7 cm/sec. Ripple measurements were made when the animals were moving ≤ 0.5 cm/sec.

Following recordings, neurons for single-unit recordings were sorted into individual units by tetrode mode-based cluster analysis in feature space using Offline Sorter software (Plexon Inc., Dallas, TX). Autocorrelation and crosscorrelation functions of spike times were used as separation tools. Only units with clear refractory periods and well-defined cluster boundaries were included in the analyses. Pyramidal cells and interneurons were distinguished based on autocorrelation plots (peak within 10 msec representing bursting), waveforms (broad waveforms, with a peak to valley spike width of >300 μsec) and mean firing rates (<5 Hz during baseline recording)^56^. Only pyramidal cells were included in analyses.

For simultaneous optogenetic stimulation and LFP recording, fiber optic probes (200 μm diameter) were connected to a Plexbright 4-channel optogenetics controller through a wired tether, and Radiant software (Plexon, Inc.) was used to drive light stimulation. Electrophysiological recordings were made from awake, behaving mice during periods of run and rest (behavioral state not separated for these experiments) using 32 channel head stages digitized at 16-bit resolution and acquired at 40 kHz using the OmniPlex D Neural Data Acquisition System (Plexon, Inc.). Continuous neural data were low pass filtered at 500 Hz and sampled at 1000 Hz. For these experiments, baseline recordings were obtained for several minutes before delivering fiber optic stimulation. Stimulation consisted of 5 pulses delivered at 10 Hz, with each pulse being 5 msec in duration and with a current intensity of 200 mA (corresponding to 10-15 mW) delivered to the light emitting diode. One train was delivered per minute for 3 minutes. Animals were then administered either vehicle or CNO (5 mg/kg, SQ), and LFP responses to light stimulation were made following identical stimulation parameters between 20 minutes and 24 hours following vehicle/CNO treatment. Response amplitudes were measured from evoked voltage deflections time locked to the optogenetic stimulation events.

#### Electrode localization

Upon completion of electrophysiology studies, mice were perfused with 4% paraformaldehyde. Heads with electrodes remaining in place in brains were then submerged in 4% paraformaldehyde for 24-48 h. Electrodes were carefully removed and brains were submerged in 30% sucrose/PBS and sectioned at 40 μm on a cryostat or vibratome.

#### Electrophysiology Data Analysis

The experimenter was blind to the genotype of animals at the time of data analysis. All neuronal data analyses were performed using Neuroexplorer software (Nex Technologies, TX) and Matlab (MathWorks, Inc., Natick, MA) with the Chronux toolbox for Matlab (http://chronux.org/). Statistical analyses were performed using GraphPad Prism version 6.

Identical analyses were used for all hM3Dq and hM4Di spectral measures. Data were first divided into periods of running (>7 cm/sec) or resting (<0.5 cm/sec and limited to up to 20 sec once an animal has started moving <0.5 cm/sec). These LFP subsets were then z-scored to control for changes in overall signal amplitude (and, consequently, power) over the course of up to 2 weeks of recordings (in the case of hM3Dq animals in which multiple doses of CNO or vehicle were administered every 2 to 3 days). LFPs were then filtered using a zero-phase offset filter in the theta (5-10 Hz), beta (14-18 Hz), low gamma (30-60 Hz) or high gamma (65-100 Hz) range. The Chronux function mtspectrum, a multitaper spectral estimate, was used with 5 tapers, and resulting spectral values were smoothed. For all treatments, spectral measures were made during each of run and rest periods during the 30 to 60 minutes following treatment. Spectral density plots for each behavioral state, each treatment and in each recording site were averaged across animals according to genotype and AAV infused. Peak powers in each frequency range were collected to compare changes in peak theta, beta, low gamma or high gamma power according to treatment. Cross frequency coupling of theta phase and low gamma (30-55 Hz) power were also measured from hippocampal LFPs during periods of running following each treatment using the method of ^57,58^. Coherence measures were performed using the Chronux function cohgramc^59^, and mean low gamma coherence was measured over the 30-60 Hz frequency range from the run and rest subsets described above. Power and coherence values measured for each treatment were compared using appropriate statistical tests, listed in text, after data were checked for normal distributions and equal variance.

Ripple events were identified according to modified methods previously described^9,37^ from recordings originating from the pyramidal cell layer during periods of rest in hM3Dq (8 animals with dorsal CA1 recordings) and hM4Di (2 animals with dorsal CA1 recordings, 4 animals with intermediate CA1 recordings) animals. These signals were denoised with an IIR notch filter at 60 and 180 Hz and filtered between 100 and 300 Hz with a 69-order FIR zero phase shift filter. Signals were then Hilbert transformed, and the absolute value envelopes were smoothed with a 50-msec window. Envelope amplitude deflections that exceeded 3 standard deviations from the mean amplitude (i.e., mean +3 standard deviations) for more than 30 msec were counted as ripple events. Deflections within 200 msec of a previous ripple event excluded. Ripple event frequency and ripple amplitude were measured and appropriate statistical tests were applied, as listed in the text, after data were checked for normal distributions and equal variance.

#### Behavioral measures

For all behavioral measures, the experimenter was blind to the genotype of animals. Estrous cycle was not tracked in female mice. All behavioral assays were performed during the light cycle, between 8 AM and 3 PM.

Two cohorts of *Amigo2*-icreERT2 mice were tested for behavioral analyses. One cohort was evaluated in assays for social approach, and another cohort was evaluated in assays for prepulse inhibition of acoustic startle responses and spontaneous alternation in a Y-maze. Between the time of performing the acoustic startle response and spontaneous alternation assay, *Amigo2*-icreERT2+ and *Amigo2*-icreERT2-animals were removed from the group for histological confirmation of hM4Di expression. For all assays, data were checked for normal distribution and equal variance before choosing a statistical test. Estrous cycle was not tracked in female mice. All behavioral assays were performed during the light cycle, between 8 AM and 3 PM.

CNO treatment for behavioral assays: hM4Di AAV-infused *Amigo2*-icreER mice undergoing behavioral studies were treated with 5 mg/kg CNO, IP, at specific times stated below.

Social approach in a three-chamber choice test: CNO was administered 20 minutes before the beginning of the three-chamber test. Each session consisted of two ten-minute phases: a habituation period and a test for sociability. For the sociability assay, mice were given a choice between visiting a side of the chamber with a novel sex-matched conspecific and visiting a side with no conspecific in it. The social testing apparatus was a rectangular, three-chambered box fabricated from clear Plexiglas (schematized in Fig. 7A). Dividing walls had doorways allowed access into each chamber. An automated image tracking system (Noldus Ethovision) provided measures of entries and duration in each side of the social test box as well as time in spent within 5 cm of the Plexiglass cages (the cage proximity zone). Times on each side are presented.

At the start of the test, the test mouse was placed in the middle chamber and allowed to explore for ten minutes, with the doorways into the two side chambers open. Measures of the amount of time spent in each side chamber were taken. After this habituation period, the test mouse was enclosed in the center compartment of the social test box, and an unfamiliar C57BL/6J “stranger” was placed in one of the side chambers. The unfamiliar mouse was enclosed in a small Plexiglass cage drilled with holes, which allowed nose contact, but prevented fighting. An identical empty Plexiglass cage was placed in the opposite side of the chamber. Following placement of the unfamiliar mouse in the empty cage, the doors were re-opened, and the subject was allowed to explore the entire social test box for a ten-minute session.

Prepulse Inhibition: Prepulse inhibition of acoustic startle responses, an index of sensorimotor gating, was measured using an SR-LAB system (San Diego Instruments, San Diego, CA). hM4Di AAV-infused animals were administered CNO 30 minutes before beginning the prepulse inhibition assay. Mice were placed inside the animal enclosure fitted to the size of the animal. The sound proof chamber was then closed and the 20-min session begun. Background white noise at 70 dB occurred throughout the session except during pre-pulse and pulse stimuli. Pre-pulse stimuli were 73, 76 or 82 dB, 20 msec long, and presented 100 msec before the pulse, which was 120 dB and 40 msec long. Movements initiated by pulse stimuli were transduced into startle amplitude. A session started with a 5-min acclimation period followed by three consecutive blocks of trials. The first and last blocks consisted of six pulse-alone trials. The middle block consisted of 52 pulse alone, prepulse+pulse, and no-stimulus (background white noise only) trials in a pseudo-randomized order. The no-stimulus trials were incorporated to measure basal movement of the animal. Each trial was separated by a random, variable 15-sec inter-trial interval. Each trial started with a 50 msec null period and ended with a 200 msec recording period.

Spontaneous alternation: hM4Di AAV-infused mice were tested in the spontaneous alternation task using a Y maze (Med Associates, consisting of three 36.83 × 7.5 cm runways covered with removable clear Plexiglass). Red lights at 3-5 lux illuminated the room, and visual cues were positioned on all four walls at a height above the maze. At the start of a trial, the Plexiglass was removed from the top of one arm of the Y-maze, a mouse was placed on a distal end of the arm, and the Plexiglass was replaced. After a 10-sec delay, the door separating the arm from the rest of the maze was opened, and the mouse was allowed to explore the maze freely for 8 min. Trials were videotaped, and an experimenter blind to the genotype scored entries of the mouse into arms during the 8-min trial. Entry into an arm required all four limbs to be within the arm. An alternation was scored when an animal entered three different arms sequentially. Percent spontaneous alternation was calculated by dividing the number of alternations by the maximum possible number of alternations as follows: (# of alternations) / (total number of arm entries – 2).

#### Results Reporting and Data Availability

For each experiment presented within the Results section and in figures, the number of replicates is presented as “N” when indicating the number of animals that were used for the experiment or as “n” when referring to the number of neurons used for the experiment. Statistical tests used for each experiment are presented in the text. Statistical significance was based on a p-value of 0.05. All error bars in graphs represent standard error of the mean. The data supporting the findings of this study are available from the corresponding author upon request.

## CONTRIBUTIONS

G.M.A., L.Y.B., S.F., D.L., N.V.R., S.S.M., and S.M.D. conceived of and designed the studies. G.M.A., L.Y.B., S.F., D.L., C.P., N.V.R., and P.J. conducted experiments and analyzed data. B.G. and N.W.P. generated transgenic mice. G.M.A and S.M.D. wrote the manuscript. S.S.M., P.J., and S.M.D. supervised the project.

## ACKNOWLEDGEMENTS

This research is supported by the Intramural Research Program of the U.S. NIH, National Institute of Environmental Health Sciences (Z01 ES100221 to S.M.D.). The UNC Mouse Behavioral Phenotyping Laboratory is supported by U.S. NIH, National Institute of Child Health and Human Development (U54HD079124 to Joseph Piven). We wish to thank Guohong Cui, Jesse Cushman and the members of the Dudek lab for critically reading this manuscript, as well as Jesse Cushman and the NIEHS Neurobehavioral Core, the Fluorescence Microscopy and Imaging Center, and the animal care staff at NIEHS for their support.

